# In vivo reprogramming leads to premature death due to hepatic and intestinal failure

**DOI:** 10.1101/2022.05.27.493700

**Authors:** Alberto Parras, Alba Vílchez-Acosta, Gabriela Desdín-Micó, Calida Mrabti, Cheyenne Rechsteiner, Fabrice Battiston, Clémence Branchina, Kevin Pérez, Christine Sempoux, Alejandro Ocampo

## Abstract

**SUMMARY:** The induction of cellular reprogramming by forced expression of the transcription factors OCT4, SOX2, KLF4, and C-MYC (OSKM) has been shown to allow the dedifferentiation of somatic cells and ameliorate age-associated phenotypes in multiple tissues and organs. Yet to date, the benefits of in vivo reprogramming are limited by the occurrence of detrimental side-effects. Here, using complementary genetic approaches, we demonstrated that continuous in vivo induction of the reprogramming factors leads to hepatic and intestinal dysfunction resulting in decreased body weight and premature death. By generating a novel transgenic reprogrammable mouse strain, which avoids OSKM expression in both liver and intestine, we drastically reduced the early lethality and adverse effects associated with in vivo reprogramming. This new reprogramming mouse allows safe and long-term continuous induction of OSKM and might enable a better understanding of in vivo reprogramming as well as maximize its potential effects on rejuvenation and regeneration.

## INTRODUCTION

The simultaneous forced expression of four transcription factors (4F), OCT4, SOX2, KLF4, and MYC (OSKM) has been shown to be sufficient to activate pluripotency networks and dedifferentiate somatic cells to a pluripotent state (Takahashi and Yamanaka, 2006). Interestingly, it has been demonstrated that transient induction of reprogramming can reverse age-associated features in cells in vitro (Lapasset et al., 2011; Olova et al., 2019; Roux et al., 2021; Sarkar et al., 2020), suggesting its capacity to alter both cellular identity and age. Likewise, the conversion of differentiated cells into pluripotency has been confirmed in vivo by multiple groups including ours (Abad et al., 2013; Ocampo et al., 2016), opening the door for potential in vivo rejuvenation and regeneration of tissues and organs. In this line, in vivo reprogramming can induce beneficial effects on skin (Doeser et al., 2018), heart (Chen et al., 2021), skeletal muscle (de Lazaro et al., 2019; Wang et al., 2021) and liver (Hishida et al., 2022) regeneration. Furthermore, reprogramming of aged mice has the capacity to improve memory (Rodriguez-Matellan et al., 2020), regenerate skeletal muscle and pancreas following injury (Chiche et al., 2017; Ocampo et al., 2016), restore vision loss in injured retina (Lu et al., 2020), and recently, rejuvenate multiple tissues and organs (Browder et al., 2022; Chondronasiou et al., 2022). Finally, cyclic induction of reprogramming has been shown to extend lifespan in a progeria mouse model (Alle et al., 2021; Ocampo et al., 2016). All these observations represent a proof of concept and highlight the potential of in vivo reprogramming for the improvement of human healthspan. However, despite many benefits, previous studies have also indicated that continuous expression of OSKM in vivo can lead to cancer development and teratoma formation (Abad et al., 2013; Mosteiro et al., 2016; Ohnishi et al., 2014), and most importantly, early mortality (Abad et al., 2013; Hishida et al., 2022; Ocampo et al., 2016; Rodriguez-Matellan et al., 2020). Consequently, these strong adverse effects represent a barrier for the safe and long-term continuous induction of reprogramming in vivo, and therefore limit its potential benefits. Importantly, the etiology of the side-effects associated with in vivo reprogramming remains unexplored, and the ultimate cause of death of reprogrammable mice is still unclear.

In order to gain insight into the adverse effects of in vivo reprogramming, we performed a comparative analysis of different reprogrammable mouse strains upon induction of OSKM. As expected, continuous expression of reprogramming factors is highly toxic, indicated by decreased body weight, temperature loss, decreased activity, and ultimately mortality. Using genetic strategies to activate or inactivate the expression of the reprogramming factors in hepatocytes and intestinal epithelial cells, we first demonstrated that specific expression of OSKM in the liver and intestine compromises the normal function of these organs and is lethal. Importantly, these severe adverse phenotypes were drastically reduced in a novel reprogrammable mouse strain where OSKM expression was avoided in the liver and intestine. Finally, we showed that safe and long-term continuous induction of in vivo reprogramming can be achieved in the absence of detrimental side effects by avoiding the expression of the Yamanaka factors in these organs. We anticipate that these observations and this novel mouse strain might open the door for reaching the maximum beneficial potential of in vivo reprogramming.

## RESULTS

### In vivo reprogramming results in hepatic and intestinal dysfunction

Despite the potential beneficial effects of in vivo reprogramming, the induction of OSKM at the organismal level has been associated with body weight loss, tumor development, and early mortality (Abad et al., 2013; Mosteiro et al., 2016; Ohnishi et al., 2014). Importantly, although certain short-term induction protocols, which allow reprogrammable mice to survive, lead to the development of tumor and teratomas, the timing of these events is not compatible with the early mortality observed upon continuous induction of in vivo reprogramming. For these reasons, in order to better understand the deleterious consequences of this process, we first performed side-by-side comparisons of two of the most studied reprogrammable mouse strains, which have been previously used to investigate the effects of in vivo reprogramming at tissue and organismal levels. These strains included the 4Fj developed by the laboratory of Rudolf Jaenisch (Carey et al., 2010) and 4Fs-B generated by the group of Manuel Serrano (Abad et al., 2013). Importantly, these strains are similar but not identical. Although each of them carries the transcriptional activator (rtTA-M2) within the ubiquitously expressed *Rosa26* locus, the doxycycline-inducible polycistronic cassette encoding the four murine factors *Oct4* (*Pou5f1*), *Sox2, Klf4* and *c-Myc* (TetO 4F) is inserted in different genomic locations, specifically in the *Col1a1* (4Fj) and *Pparg* (4Fs-B) loci (Figure 1A). First, to induce the expression of the reprogramming factors in vivo, 4Fj and 4Fs-B mice were treated with doxycycline in drinking water (1 mg/ml) continuously (Figure S1A). As expected, both lines experienced body weight loss (Figure 1B), albeit to different extent, lower activity (Figure 1C), and decreased body temperature (Figure S1B) after only a few days of treatment, indicating early toxicity of in vivo reprogramming. Unsurprisingly, mice of both strains began to die after 3 days of doxycycline treatment, with a significantly different median survival of 5 days and 10 days for the 4Fj and 4Fs-B respectively (Figure 1D). Interestingly, gross postmortem examination of these animals showed neither indication of tumor nor teratoma formation, suggesting that these events were not the cause of premature death. For this reason, in order to investigate the early and differential mortality associated with the expression of the Yamanaka factors in vivo, we analyzed the expression of the reprogramming factors in multiple tissues and organs. Although *Oct4* and *Sox2* transcripts were detected at similar levels in both strains in organs such as small intestine and kidney, a higher expression of these factors was observed in both liver and blood of 4Fj compared to 4Fs-B mice (Figure 1E and Figure S1C). In the same line, OCT4 was detected in the small intestine of both strains at the protein level, but only in the liver of 4Fj animals (Figures S1D). Notably, we observed a significant increase of Ki67 positive cells in the liver of the 4Fj mice as well as the intestine of both strains, indicating that reprogramming can induce cellular proliferation in these organs (Figure S1F) in agreement with previous reports (Hishida et al., 2022; Ohnishi et al., 2014).

**Figure 1:**
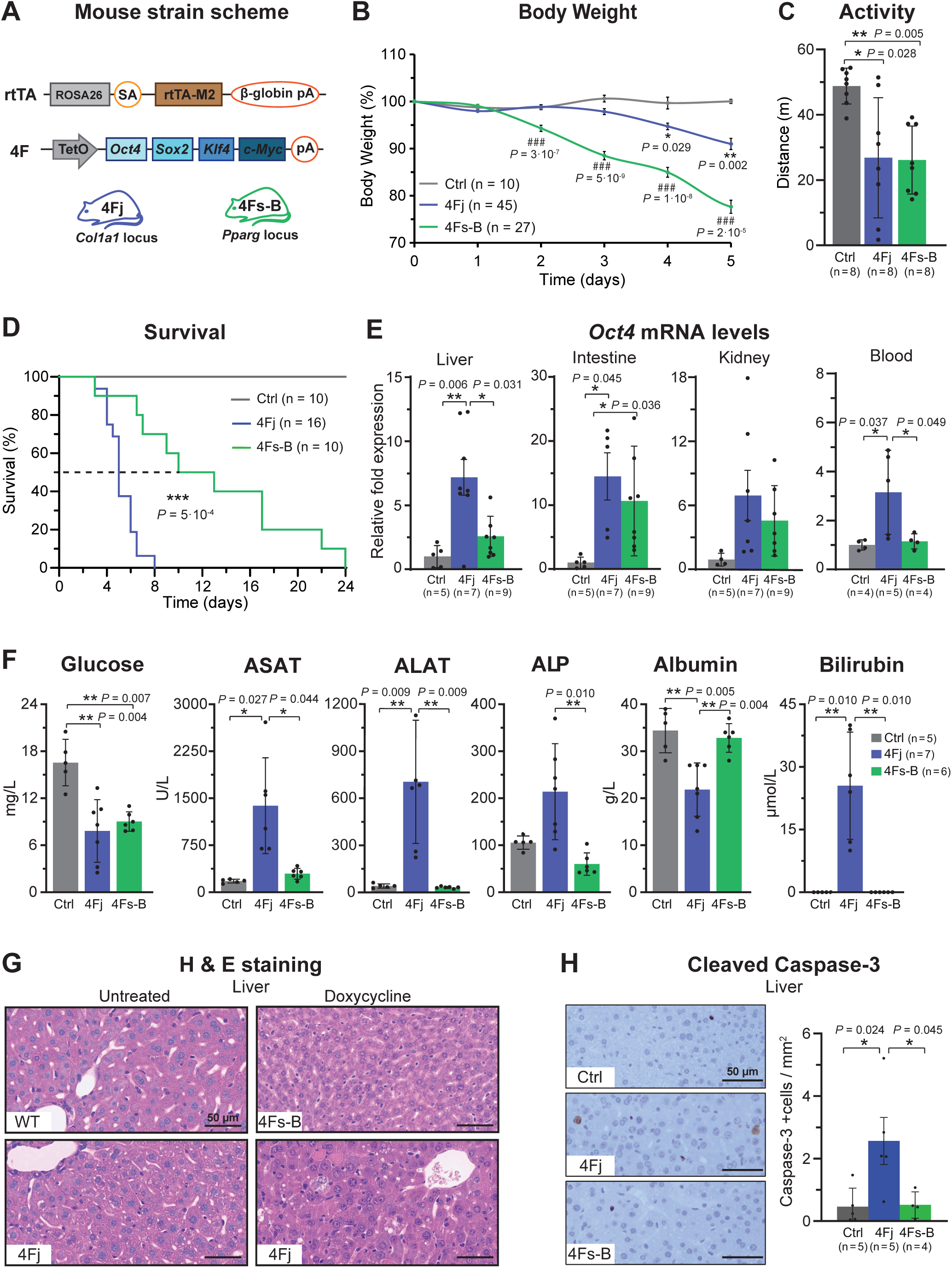
In vivo reprogramming results in hepatic and intestinal dysfunction in different reprogrammable mouse strains. (A) Schematic representation of reprogrammable mouse strains used in this study carrying the rtTA (reverse tetracycline-controlled transactivator) transgene at the *Rosa26* locus and a doxycycline-inducible polycistronic cassette for the expression of the mouse 4F (*Oct4, Sox2, Klf4* and *c-Myc*) in the *Col1a1* locus (4Fj) and *Pparg* locus (4Fs-B), (pA) polyA sequence, (TetO) tetracycline operator minimal promoter. (B) Body weight changes in 4Fj and 4Fs-B mice upon continuous administration of doxycycline, and untreated 4F control mice (Ctrl). (C) Distance travelled in open field cage following 3.5 days of doxycycline treatment of controls, 4Fj and 4Fs-B mice. (D) Survival of 4Fj and 4Fs-B mice upon continuous administration of doxycycline and untreated 4F mice. (E) *Oct4* mRNA levels in liver, small intestine, kidney and blood in untreated controls, 4Fj and 4Fs-B mice after 4 days of doxycycline treatment. (F) Glucose, liver enzymes (ASAT, ALAT, ALP), bilirubin and albumin levels in plasma of untreated animals (Ctrl) and 4Fj and 4Fs-B mice after 4 days of induction of in vivo reprogramming. (G) Liver hematoxylin and eosin stain for an untreated wildtype and 4Fj mice, and treated 4Fj and 4Fs-B mice during 4 days. (H) Immunostaining and quantification of cleaved caspase-3 positive cells in liver of untreated controls, 4Fj and 4Fs-B mice. Data are mean ± SEM. Statistical significance was assessed by (B-C, F-G) one-way ANOVA followed by Tukey’s post hoc or Games Howell tests and (D) log-rank (Mantel-Cox) test.

Next, with the goal of investigating the potential organ dysfunction leading to the early lethality, we performed chemistry analysis of plasma collected from 4Fj and 4Fs-B mice after 4 days of continuous doxycycline treatment, the time point at which both strains were found sick. Interestingly, a significant decrease in glucose levels was observed in both strains (Figure 1F). Importantly, elevated levels of the circulating liver enzymes aspartate aminotransferase (ASAT), alanine aminotransferase (ALAT), alkaline phosphatase (ALP), and bilirubin, together with a decrease in albumin, was observed in the 4Fj but not in 4Fs-B mice, suggesting hepatic failure only in this strain (Figure 1F). In this line, analysis of 4Fj liver histology after doxycycline treatment showed a disorganized lobule architecture, with many apoptotic bodies, scattered vacuolated hepatocytes and anisonucleosis and numerous Kupffer cells. However, the liver parenchyma of treated 4Fs-B mice only showed some small droplets in the cytoplasm of the hepatocytes located in zone 2 and 3 (Figure 1G). These observations correlate with an increase in cleaved caspase-3 positive cells, a marker of apoptosis, in only 4Fj mice, confirming the hepatic damage after reprogramming in this strain (Figure 1H). Histological analysis of the small intestine showed alterations in both strains after doxycycline treatment. Specifically, the intestine of 4Fs-B mice presented shortened villi, with less goblet cells, and dedifferentiated enterocytes. In 4Fj mice, the intestine was completely atrophic, with very short villi, apoptotic figures in the crypts and dedifferentiated smaller enterocytes with complete loss of goblet cells and destruction of the brush border (Figure S1E). Lastly, no changes in potassium, phosphate, or uric acid levels were detected in neither of the strains suggesting normal kidney function at least after 4 days of in vivo reprogramming (Figure S1F).

Strikingly, our findings indicate that the adverse phenotypes associated with in vivo reprogramming can be related to organ failure prior to tumor or teratoma formation, which may indeed represent the main cause of the early mortality observed in whole body reprogrammable mouse strains.

### Liver and intestine-specific reprogramming leads to body weight loss and early mortality

Next, in order to determine whether hepatic and intestinal dysfunction resulting from in vivo reprogramming is a direct effect of the expression of the Yamanaka factors in these organs, and consequently the cause of premature death, we generated two novel tissue-specific reprogrammable mouse strains for the specific expression of reprogramming factors in these organs. Towards this goal, we used the 4Fj mice, which is the only strain characterized by high OSKM expression in both organs. For this purpose, the reverse tetracycline trans-activator (rtTA-M2) was substituted by a LoxP-STOP-LoxP (LSL) cassette-blocked rtTA trans-activator and the resulting offspring were crossed with both an Albumin-Cre mice (Postic et al., 1999) to generate the 4F Liver mice or with a Villin-Cre mice (Madison et al., 2002) to generate 4F Intestine mice (Figure 2A). As expected, the Cre lines showed expression of the Cre recombinase specifically in the liver or the intestine (Figure S2A). Moreover, we confirmed the expression of the reprogramming factors in the targeted organs upon doxycycline treatment (Figure 2B and Figure S2B). Interestingly, significant body weight loss was observed after only 2 days of doxycycline treatment in 4F Intestine mice, and 24 hours later in 4F Liver mice (Figure 2C), as well as decreased body temperature (Figure S2C). Interestingly, plasma analysis revealed liver failure associated with high levels of circulating hepatic enzymes (ASAT, ALAT and ALP) and bilirubin, as well as low albumin levels in 4F Liver mice (Figure 2E), in agreement with a recent study (Hishida et al., 2022). Conversely, we did not observe changes in liver enzymes and metabolites in 4F Intestine mice, suggesting normal liver function (Figure 2E). Instead, 4F Intestine mice displayed hallmarks of nutrient malabsorption, including a significant decrease in glucose levels in plasma (Figure 2D) and diarrhea. More importantly, all mice of both strains were found extremely sick after several days of reprogramming and succumbed to illness with median survival of 3.5 days (4F Liver) or 5 days (4F Intestine mice) (Figure 2F).

**Figure 2:**
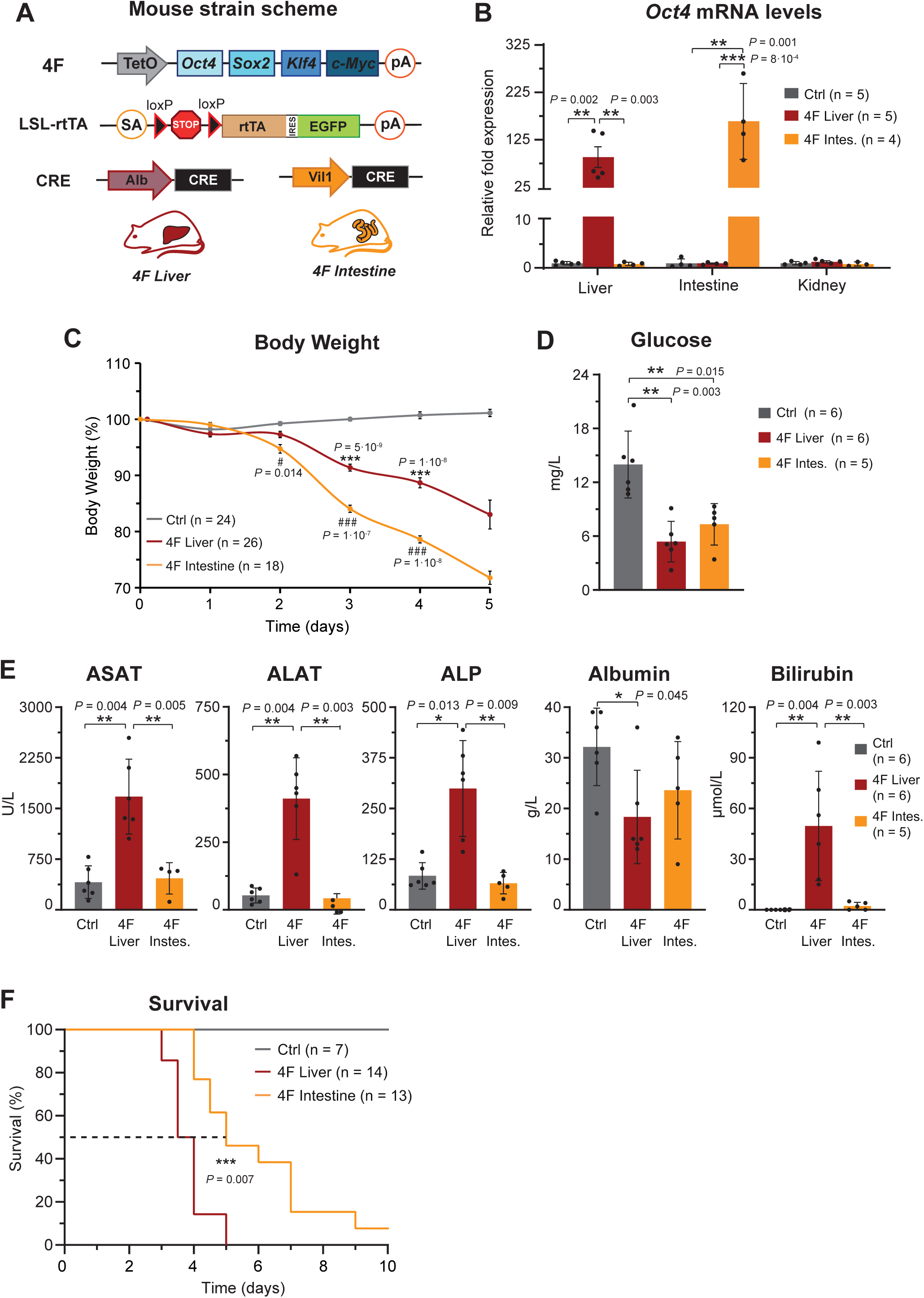
Liver or intestine-specific reprogramming leads to body weight loss and early mortality. (A) Schematic representation of liver- and intestine-specific reprogramming strains carrying a polycistronic for the expression of the mouse reprogramming factors (*OSKM*) in the *Col1a1* locus (4Fj), the LoxP-STOP-LoxP (LSL) cassette-blocked rtTA trans-activator, and Cre recombinase transgenes under the control of the promoter of the mouse *Albumin* and *Villin 1* genes. (B) *Oct4* transcript levels in the liver, small intestine, and kidney of 4F-Liver, 4F-Intestine, and control mice (carrying only the 4Fj and LSLr-tTA transgenes but no Cre) after 4 days of induction. (C) Changes in body weight of 4F-Liver, 4F-Intestine and control mice upon continuous administration of doxycycline. (D-E) Glucose, liver enzymes (ASAT, ALAT, ALP), bilirubin and albumin levels in plasma of control, 4F-Liver, and 4F-Intestine mice after 4 days of induction of in vivo reprogramming. (F) Survival of control, 4F-Liver and 4F-Intestine mice upon continuous administration of doxycycline. Data are mean ± SEM. Statistical significance was assessed by (B-E) one-way ANOVA followed by Tukey’s post hoc test and (F) log-rank (Mantel-Cox) test.

Overall, these results indicate direct negative effects of the expression of the reprogramming factors in the liver and intestine, which are sufficient to induce rapid body weight loss and ultimately lead to death. Moreover, a comparison of the phenotypes observed in these organ-specific strains and whole-body reprogrammable mice strongly suggests that most of the early negative consequences associated with in vivo reprogramming are due to hepatic and intestinal dysfunction.

### Bypassing the expression of OSKM in liver and intestine significantly reduces the adverse effects of in vivo reprogramming

To further validate that hepatic and intestinal reprogramming are major causes of early mortality in whole-body 4F mice, we generated two novel mouse strains which express OSKM in whole body with the exception of the liver or the intestine. To achieve this goal, we substituted the 4F polycistronic cassette in the 4Fj rtTA-M2 mice, by a 4Fj cassette flanked by loxP sites, which allows its Cre-mediated excision. Subsequently, we crossed these mice to Albumin-Cre or Villin-Cre in order to remove the 4F cassette in the targeted organs and generate 4F Non-Liver or 4F Non-Intestine mice respectively (Figure 3A). Next, we confirmed the decrease expression of *Oct4* and *Sox2* transcript levels in the liver or intestine of 4F Non-Liver and 4F Non-Intestine mice (Figure 3B and Figure S3A). As expected, no reduction of OSKM expression was observed in other organs such as the kidney upon doxycycline treatment when compared with whole-body 4F-Flox controls (Figure 3B and Figure S3A).

**Figure 3:**
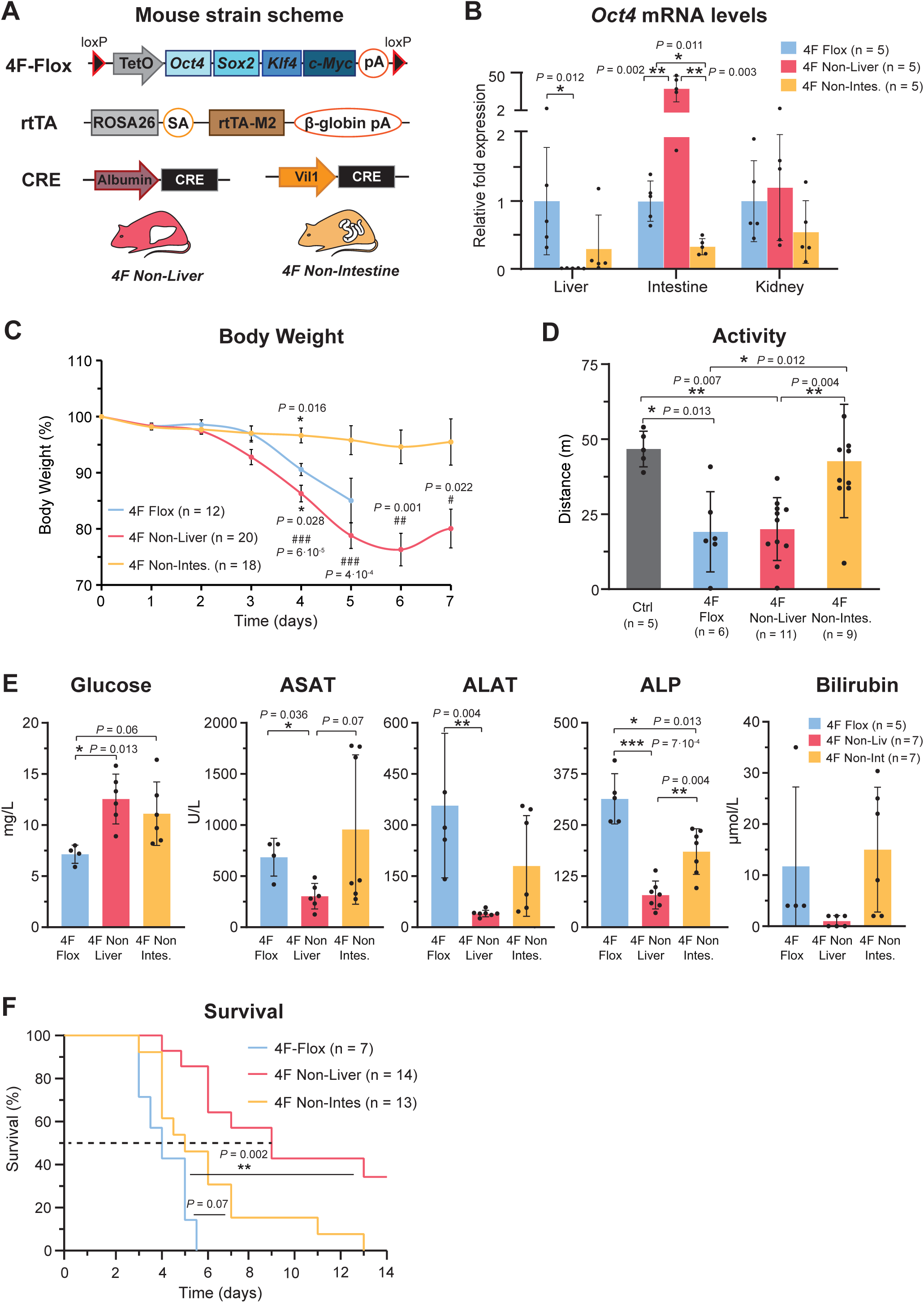
Reduction of adverse effects of in vivo reprogramming by abolishing the expression of OSKM in the liver and intestine of reprogrammable mice. (A) Schematic representation of the 4F Non-Liver and 4F Non-Intestine mouse strains carrying the polycistronic cassette for the expression of the reprogramming factors (*Oct4, Sox2, Klf4* and *c-Myc*) under a doxycycline-inducible promoter flanked by loxP sites in the *Col1a1* locus (4Fj-Flox), the reverse tetracycline-controlled transactivator (rtTA) in the *Rosa26* locus, and Cre recombinase transgenes under the control of the promoter of the mouse *Albumin* and *Villin 1* mouse genes. (B) mRNA levels of *Oct4* in the liver, small intestine, and kidney, of the 4F-Flox (whole body reprogrammable mice), 4F Non-Liver, 4F Non-Intestine mice, after 4 days of induction of in vivo reprogramming. (C) Changes in body weight of 4F-Flox (whole body reprogramming), 4F Non-Liver, 4F Non-Intestine mice, upon continuous administration of doxycycline and (D) distance travelled in open field cage following 3.5 days of treatment. (E) Glucose levels, liver enzymes (ASAT, ALAT, ALP) and bilirubin levels in plasma of 4F-Flox, 4F Non-Liver, 4F Non-Intestine after 4 days of reprogramming. (F) Survival of 4F-Flox (whole body), 4F Non-Liver and 4F Non-Intestine mice upon continuous administration of doxycycline (1mg/ml). Data are mean ± SEM. Statistical significance was assessed by (B-E) one-way ANOVA followed by Tukey’s post hoc test and (F) log-rank (Mantel-Cox) test.

Importantly, 4F Non-Intestine mice did not manifest abnormalities in body weight loss or activity upon induction of in vivo reprogramming (Figure 3C-D). Moreover, none of the mice of this line showed diarrhea or changes in the glucose level in plasma, suggesting that intestinal function was not affected (Figure 3E). On the other hand, ASAT, ALAT, ALP and bilirubin levels were highly elevated, indicating hepatic failure after 4 days due to hepatic reprogramming (Figure 3E). More importantly, none of the animals survived the continuous treatment with doxycycline treatment, showing a median survival comparable to 4F-Flox controls of only 5 days (Figure 3F).

Conversely, 4F Non-Liver mice did not show any sign of liver failure, as observed by the low levels of ASAT, ALAT ALP and bilirubin in the plasma (Figure 3E). However, despite normal liver function, 4F Non-Liver mice displayed decreased activity and weight loss starting after 3 days of treatment probably due to intestinal reprogramming (Figure 3C-D). Remarkably, the median survival in this strain was significantly extended to 9 days (Figure 3F).

Taken together, these data suggest that intestinal reprogramming leading to nutrient malabsorption is one of the main causes of early adverse effects of reprogramming, including body weight loss, diarrhea and low activity, whereas hepatic reprogramming triggers liver failure and represents the primary cause of premature death in 4Fj reprogrammable mouse strain. Moreover, avoiding the expression of OSKM in these organs partially prevents the adverse effects of in vivo reprogramming including decrease in body weight and premature death.

### Safe and long-term continuous reprogramming can be achieved in the absence of hepatic and intestinal expression of OSKM

Finally, with the goal of achieving the safe and long-term induction of in vivo reprogramming, we generated a novel quadruple transgenic mouse strain in which the 4F cassette (4F-Flox) is simultaneously removed from both liver and intestine (4F Non-Liver/Intestine mice) (Figure 4A). In order to validate this novel strain, we first confirmed the absence of OSKM expression in the liver and intestine of these mice 4 days after doxycycline treatment (Figure 4B and Figure S4A). Surprisingly, none of the adverse effects associated with in vivo reprograming including decrease in body temperature (Figure 4SB), diarrhea, body weight loss, or abnormal activity (Figure 4C-D) were detected in this strain following the induction by doxycycline treatment. Moreover, no alterations in plasma parameters characteristic of liver failure, such as, albumin, ASAT, ALAT, or ALP were detected (Figure 4E). Interestingly and compared to the whole-body 4F-Flox, the first 4F Non-Liver/Intestine mouse died after more than one week of continuous induction of reprogramming, and 60% of these mice survived up to one month of continuous doxycycline treatment, making this strain, the reprogrammable line known to date that can survive the longest protocols of continuous induction of in vivo reprogramming at high dose (Figure 4F). Subsequently, 4F Non-liver/intestine mice were euthanized after a month of doxycycline treatment and multiples organs were collected. Afterwards, we confirmed that the lack of expression of the reprogramming factors in the liver and small intestine (Figure 4G and Figure S4C), and no alterations in plasma parameters of 4F Non-Liver/Intestine mice after 4 weeks of treatment, when compared with whole body reprogrammable mice after only 4 days (Figure S4D). On the other hand, comparable or even higher level of *Oct4* and *Sox2* expression were detected in kidney and blood of 4F Non-Liver/Intestine mice after continuous induction (Figure 4G and Figure S4C).

**Figure 4:**
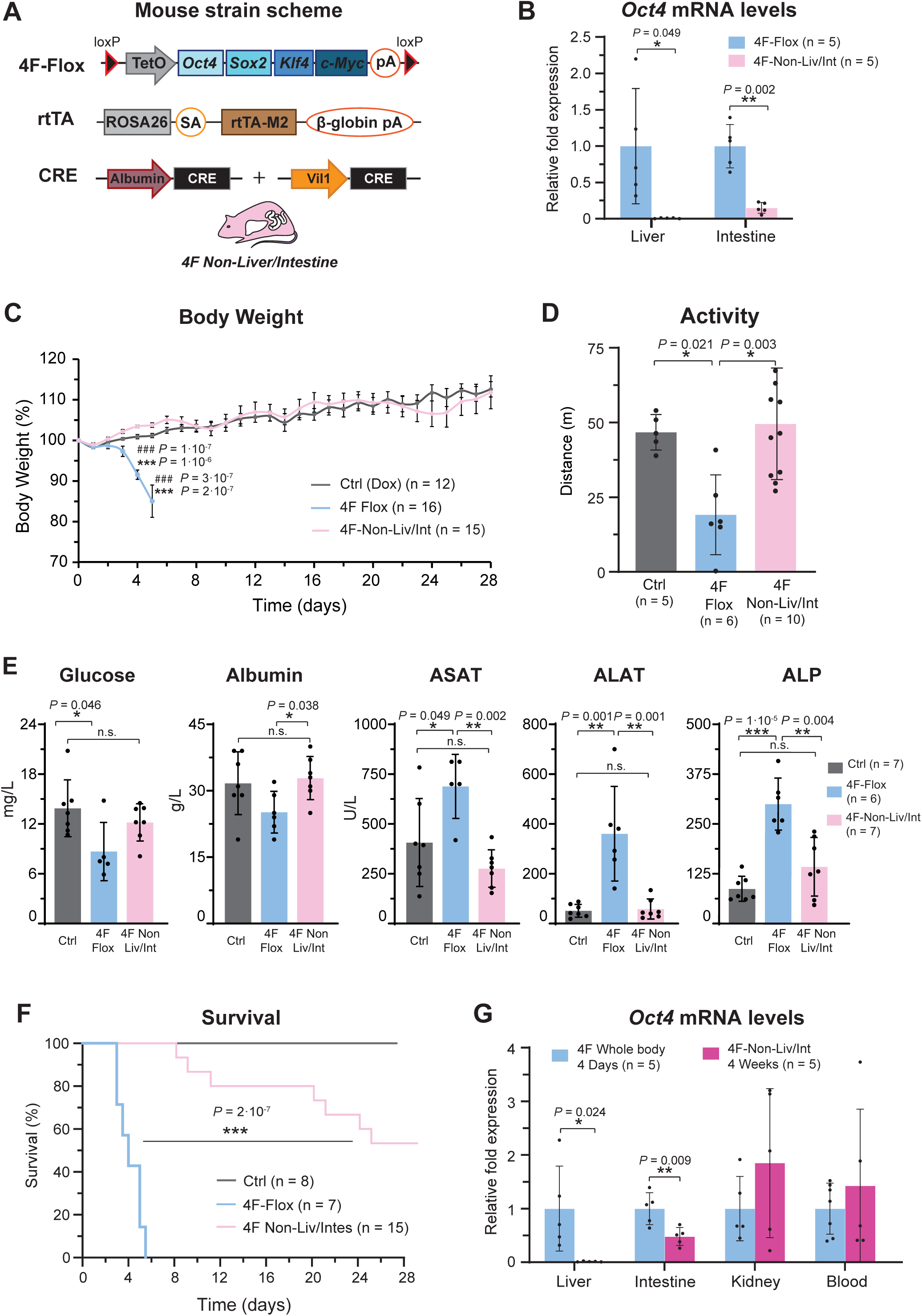
Generation of novel reprogrammable mouse strain for the safe and long-term induction of in vivo reprogramming. (A) Schematic representation the 4F Non-Liver/Intestine quadruple transgenic mouse strain carrying the polycistronic cassette for the expression of the mouse reprogramming factors (OSKM) under a doxycycline-inducible promoter flanked by LoxP sites in the *Col1a1* locus (4Fj-Flox), the reverse tetracycline-controlled transactivator (rtTA) in *Rosa26* locus and Cre recombinase under the control of the promoter of the mouse *Albumin* and *Villin 1* genes. (B) mRNA levels of *Oct4* in the liver and small intestine in 4F-Flox (whole body reprogrammable mice) and 4F Non-Liver/Intestine mice after 4 days of reprogramming induction. (C) Body weight of control (no reprogrammable mice), 4F-Flox (whole body reprogramming) and 4F Non-Liver/Intestine mice upon continuous administration of doxycycline and (D) activity measured in distance travelled in open field test following 3.5 days of treatment. (E) Glucose, liver enzymes (ASAT, ALAT, ALP) and albumin levels in plasma of controls no reprogrammable mice, 4F-Flox (OSKM whole body) and 4F Non-Liver/Intestine mice after 4 days of induction of in vivo reprogramming. (F) Survival of control, 4F-Flox (whole body), and 4F Non-Liver/Intestine mice upon continuous administration of doxycycline. (G) mRNA levels of *Oct4* in multiple organs of 4F whole body reprogrammable and 4F Non-Liver/Intestine mice after 4 days and 4 weeks respectively, of continuous doxycycline treatment. Data are mean ± SEM. Statistical significance was assessed by (B, G) two-sided unpaired t-test and Mann-Whitney-Wilcoxon test, (C-E) one-way ANOVA followed by Tukey’s post hoc test and (F) log-rank (Mantel-Cox) test.

Lastly and with the goal of confirming the safety of the long-term induction of in vivo reprogramming in 4F Non-Liver/Intestine mice, we induce the expression of the reprogramming factors for 4 days or 2 weeks and monitor these mice for 2 months after doxycycline withdrawal. Importantly, none of the mice died after treatment withdrawal (Figure 4SE), none the adverse effects described above typically associated with whole-body reprogramming such us body weight loss were observed, and no visible tumors were detected in these mice neither during the doxycycline treatment or up 2 months following its withdrawal (Figure 4SF).

Taken together, these results demonstrate that avoiding OSKM expression in liver and intestine allows the continuous induction of in vivo reprogramming for a longer period of time without significant adverse effects. For these reasons, this novel reprogrammable mouse strain might open the door for enhancing and extending the beneficial effects of reprogramming to other tissues and organs.

## DISCUSSION

The transient induction of cellular reprogramming has been shown to improve regeneration in multiple organs and ameliorate age-associated features both in vitro and in vivo, opening the door to a new era of regenerative medicine and possible reversion of both molecular and physiological age-associated phenotypes. Despite its huge potential, the field of in vivo reprogramming is still in its initial stages of development with most studies being published in the past few years (Taguchi and Yamada, 2017). In contrast to the studies of cellular reprogramming in vitro, there is a profound lack of understanding of the effects of reprogramming in a living organism with many open questions still unresolved. As an example, although tumor and teratoma formation after continuous expression of OSKM has been well-reported (Abad et al., 2013; Mosteiro et al., 2016; Ohnishi et al., 2014), remains unclear whether these events or alternative mechanisms represent the major cause of premature death in reprogrammable mice. Therefore, in order to avoid the adverse effects and early lethality associated with in vivo reprogramming, most studies have typically focused on the short-term expression of the reprogramming factors at organismal level, or more recently, the tissue- or organ-specific expression. In this line, to induce whole body expression of OSKM in the absence of side effects, researchers have commonly used short-term cyclic expression of the OSKM factors, such as 2 days (Browder et al., 2022; Ocampo et al., 2016) or 3 days (Rodriguez-Matellan et al., 2020) of doxycycline (1mg/ml) followed by 5 or 4 days withdrawal, or lower concentration of doxycycline (0.2 mg/ul) for 7 days (Chondronasiou et al., 2022). Alternatively, the generation of tissue-specific reprogrammable mouse models such as skeletal muscle-specific (Wang et al., 2021), heart-specific (Chen et al., 2021; de Lázaro et al., 2021) and liver-specific (Hishida et al., 2022), or the ectopic local expression of Yamanaka factors by using adeno-associated virus (Lu et al., 2020; Senis et al., 2018) have allowed to circumvent the negative consequences of organismal reprogramming. Unfortunately, although these protocols have been shown to be effective on improving the regeneration capacity of some specific organs and ameliorating some age-associated phenotypes, it appears that the short-term or mild induction of in vivo reprogramming is insufficient to induce significant systemic rejuvenation or extend the lifespan of wild type mice. Thus, it seems that novel strategies based on better understanding of the biology of in vivo reprogramming will be necessary to achieve its induction in the absence of adverse side-effects and therefore maximize its potential.

Toward this goal, the comparative analysis of different reprogrammable strains performed in this study indicates that despite their similarities, different reprogramming strains behave differently and are characterized by differential expression of OSKM in multiple tissues and organs. Moreover, our findings demonstrate that hepatic and intestinal reprogramming directly leads to dysfunction and damage in these organs causing strong adverse phenotypes, which are lethal within a few days. Importantly, the cause of death of reprogrammable mice has been a fundamental and long-standing question even a decade after these strains were generated. We hypothesize that the sensitivity of liver and intestine to cellular reprogramming might be due to multiple factors including their role in the digestive system and their proliferative and regenerative nature. Consequently, organs implicated in the absorption and metabolism of nutrients, such as the intestinal tract, liver, and kidney, are expected to be highly exposed to doxycycline and therefore might express higher levels of Yamanaka factors. On the other hand, the intestinal epithelium is one of the faster self-renewing tissues (Tetteh et al., 2015) and adult hepatocytes exhibit spontaneous cellular reprogramming during liver regeneration (Yanger et al., 2013) indicating high plasticity and susceptibility of these cell types to reprogramming and dedifferentiation, and potentially leading to loss of function upon OSKM expression.

Based on these observations and by generating a novel reprogrammable mouse strain where expression of OSKM is avoided in the liver and intestine, we have reduced the detrimental effects of continuous induction of OSKM including early mortality, allowing safe long-term induction of in vivo reprogramming. Nevertheless, we cannot rule out the possibility that removing the expression of the factors in these organs might cancel some positive indirect effects of reprogramming at the organismal level. In this line, it is tempting to speculate that the decrease in nutrient absorption and body weight observed following the induction of intestinal reprogramming could be responsible for some of the systemic benefits.

Importantly, although this new generation of reprogramming strains allows for long-term continuous reprogramming in the whole body without early adverse effect and lethality, the fact that some mice ultimately died after extended periods indicates that continuous reprogramming was still partially toxic, maybe by affecting other tissues or organs that remain unknown. For these reasons, it would be essential to continue investigating the effects of in vivo reprogramming in additional organs and develop strategies to better control the expression of the factors in these organs.

We believe that despite significant advances in this area of research and its enormous translational potential, we have so far failed to extend to living animals many of the beneficial effects of partial reprogramming observed in cells in vitro. Thus, it is possible that recent data on the potential benefits of in vivo reprogramming, specially in wild type mice, has been somehow not as significant as expected maybe due to the use of short-term reprogramming protocols designed to avoid early adverse effects and premature death.

Toward this ultimate goal, our findings represent initial advances in the knowledge of this emergent field of research and open new avenues for the long-term and safer induction of in vivo reprograming in the absence of adverse effects, which we believe will be critical for its clinical translation. Therefore, we hope that these findings can serve as the basis for the study of organismal regeneration and rejuvenation, ultimately leading to the amelioration of diseases and improvements in human health and lifespan.

## EXPERIMENTAL PROCEDURES

### Animal housing

All the experimental procedures were performed in accordance with Swiss legislation after the approval from the local authorities (Cantonal veterinary office, Canton de Vaud, Switzerland). Animals were housed in groups of five mice per cage with a 12hr light/dark cycle between 06:00 and 18:00 in a temperature-controlled environment at around 25°C and humidity between 40 and 70% (55% in average), with free access to water and food. Wild type (WT) and transgenic mouse models used in this project were generated by breeding and maintained at the Animal Facility of Epalinges and the Animal Facility of the Department of Biomedical Science of the University of Lausanne.

### Mouse strains

All WT and transgenic mice were used in C57BL/6J background. Whole-body reprogrammable mouse strain 4Fj rtTA-M2, carrying the OSKM polycistronic cassette inserted in the *Col1a1* locus and the rtTA-M2 trans-activator in *Rosa 26* locus (rtTA-M2), was generated in the laboratory of professor Rudolf Jaenisch (Carey *et al*., 2010) and purchased from The Jackson Laboratory, Stock No: 011004. The reprogrammable mouse strain 4Fs-B rtTA-M2, carrying the OSKM polycistronic cassette inserted in the *Pparg* locus and the rtTA-M2 trans-activator in *Rosa 26* locus (rtTA-M2), were previously generated by professor Manuel Serrano (Abad *et al*., 2013) and kindly generously donated by professor Andrea Ablasser. 4F-Liver and 4F-Intestine reprogrammable mouse strains were generated by substituting the rtTA-M2 of the 4Fj with a lox-stop-lox rtTA (LSLrtTA), (Belteki et al., 2005), purchased in The Jackson Laboratory, Stock No: 005670. The resultant offspring was crossed with an Albumin-cre, Stock No 003574, (Postic *et al*., 1999) or Villin-cre, Stock No 021504, (Madison *et al*., 2002) strains expressing selectively Cre recombinase in the liver and intestine, respectively.

The 4F Non-liver and 4F Non-intestine mouse strains were generated by breeding the 4F-Flox strain, carrying loxP sites flanking the 4F cassette in the *Col1a1* locus, previously generated by professor Jaenisch (Carey *et al*., 2010), and purchased from The Jackson Laboratory, Stock No: 011001, with Albumin-Cre (4F Non-liver), Villin-Cre (4F Non-intestine) or Albumin-Cre and Villin-Cre (4F Non-Liver/Intestine) mice to specifically remove the 4F cassette in these organs. All transgenic mice carry the mutant alleles in heterozygosity.

### Doxycycline administration

In vivo expression of OSKM in all reprogrammable mouse strains was induced by continuous administration of doxycycline (Sigma, D9891) in drinking water (1 mg/ml) in 2-3-month-old mice.

### Mouse monitoring and euthanasia

All mice were monitored at least three times per week. Upon induction of in vivo reprogramming, mice were monitored daily to evaluate their activity, posture, alertness, body weight, presence of tumors or wound, and surface temperature. Body temperature was recorded using non-contact infrared thermometer (SBS-IR-360-B, Steinberg). Mice were euthanized according to the criteria established in the scoresheet. We defined lack of movement and alertness, presence of visible tumors larger than 1cm^3^ or opened wounds, body weight loss of over 30% and surface temperature lower of 34ºC as imminent death points.

For survival and body weight experiments only males were used, and for tissue and organ collection, mice of both genders were randomly assigned to control and experimental groups. Animals were sacrificed by CO_2_ inhalation (6 min, flow rate 20% volume/min). Subsequently, before perfusing the mice with saline, blood was collected from the heart. Finally, multiple organs and tissues were collected in liquid nitrogen and used for DNA, RNA, and protein extraction, or placed in 4% formaline for histological analysis.

### Mouse activity

Locomotor activity was assessed by Open Field test in 2-month-old mice after 3.5 days of continuous treatment of doxycycline. Briefly, mice were individually placed in the center of a Plexiglas boxes (45 × 45 × 38 cm, Harvard Apparatus, 76-0439). Mice movements were recorded for 15 minutes (Stoelting Europe, 60516) and then analyzed using ANY-maze video tracking software (ANY-maze V7.11, Stoeling).

### RNA extraction

Total RNA was extracted from mouse tissues and organs using TRIzol (Invitrogen, 15596018). Briefly, 500 µl of TRIzol was added to 20-30 µg of frozen tissue into a tube (Fisherbrand 2 ml 1.4 Ceramic, Cat 15555799) and homogenized at 7000 g for 1 min using a MagNA Lyser (Roche diagnostic) at 4ºC. Subsequently, 200 µl of chloroform was added to the samples and samples were vortexed for 10 sec and placed on ice for 15 min. Next, samples were centrifuged for 15 min at 12000 rpm at 4°C and supernatants were transferred into a 1.5 ml vial with 200 µl of 100% ethanol. Finally, RNA extraction was performed using the Monarch total RNA Miniprep Kit (NEB, T2010S) following the manufacture recommendations and RNA samples were stored at -80°C until use.

### cDNA synthesis

Total RNA concentration was determined using the Qubit RNA BR Assay Kit (Q10211, Thermofisher), following the manufacture instructions and a Qubit Flex Fluorometer (Thermofisher). Prior to cDNA synthesis, 2 μL of DNAse (1:3 in DNase buffer) (Biorad, 10042051) was added to 700 ng of RNA sample, and then incubated for 5 min at room temperature (RT) followed by an incubation for 5 min at 75°C to inactivate the enzyme. For cDNA synthesis, 4 μL of iScript™ gDNA Clear cDNA Synthesis (Biorad, 1725035BUN) was added to each sample, and then placed in a thermocycler (Biorad, 1861086) following the following protocol: 5 min at 25°C for priming, 20 min at 46°C for the reverse transcription, and 1 min at 95°C for enzyme inactivation. Finally, cDNA was diluted using autoclaved water at a ratio of 1:5 and stored at -20°C until use.

### Semiquantitative RT-PCR

The following specific primers were used to detect the expression of the Cre recombinase in the cDNA of mice samples, Cre forward: 5’-GAACGAAAACGCTGGTTAGC-3’, and Cre: reverse 5’-CCCGGCAAAACAGGTAGTTA-3’ at a final concentration of 0.4 µM. DNA was amplified using DreamTaq Green PCR Master Mix 2X (Thermofisher, K1081) following the amplification protocol: 3 min at 95°C + 33 cycles (30 s at 95°C + 30 s at 60 °C + 1 min at 72°C) + 5 min at 72°C. PCR product were loaded and run in an agarose (1.6%) gel containing ethidium bromide (Carlroth, 2218.1). Images were scanned with a gel imaging system (Genetic, FastGene FAS-DIGI PRO, GP-07LED).

### qRT-PCR

qRT-PCR was performed using SsoAdvanced SYBR Green Supermix (Bio-Rad, 1725274) in a PCR plate 384-well (Thermofisher, AB1384) and using a Quantstudio 12K Flex Real-time PCR System instrument (Thermofisher). Forward and reverse primers were used at a ratio 1:1 and final concentration of 5 µM with 1ul of cDNA. *Oct4* and *Sox2* mRNA levels were determined using the following primers: *Oct4* forward: 5’-GGCTTCAGACTTCGCCTTCT-3’ *Oct4* reverse: 5’-TGGAAGCTTAGCCAGGTTCG-3’, *Sox2* forward: 5’-TTTGTCCGAGACCGAGAAGC-3’, *Sox2* reverse: 5’-CTCCGGGAAGCGTGTACTTA-3’. mRNA levels were normalized using the house keeping gene *Gapdh (*forward: 5’-GGCAAATTCAACGGCACAGT-3’, reverse: 5’-GTCTCGCTCCTGGAAGATGG-3’).

### Immunohistochemistry

Mice were euthanized with CO_2_ and multiple tissues and organs were collected, placed in 4% formaline (Sigma, 252549) overnight, and then immersed in 30% sucrose in phosphate buffered saline (PBS) for 72 h. Subsequently, samples were paraffin-embedded with a Leica ASP300S tissue processor (Leica, Heerbrug, Switzerland), sections prepared with a Microm HM 335 E microtome (Thermo Scientific,Walldorf, Germany) and mounted on Superfrost Plus slides (Thermo Scientific). Next, slides were deparaffinized and rehydrated with xylol and alcohol. Each section was routinely stained with hematoxylin and eosin, mounted on glass slides, and examined. Sections were immunostained with the primary antibody for 60 min and subsequently incubated with Dako EnVision HRP secondary antibody (Dako) for 30 min. For Ki67 immunostaining, the total number of immunopositive cells was quantified in the liver in three different regions per animal and means of the three regions were calculated and normalized by area. For cleaved Caspase-3, a complete liver slice per animal was quantified and normalized by area using a Nikon microscope camera (Nikon, digital sight 1000). Antibodies: rabbit Ki67 (1:100, Abcam, ab16667); rabbit cleaved CASP3, Asp175 (1:200, Cell Signaling, 9661).

### Plasma analysis

Mouse blood was collected in lithium heparin tubes (BD microtainer, 365966), centrifugated at 6500 rpm for 5 min, and the supernatant (plasma) was transferred to a new vial. Multiple parameters (glucose, ASAT, ALAT, ALP, bilirubin, albumin, potassium, uric acid and phosphate) were analyzed and quantified by the Service de Chimie Clinique du CHUV using a Cobas 8000 (Roche Diagnostics).

### Western blot

Samples were quickly dissected on an ice-cold plate and stored at –80 °C. Protein extraction was prepared by homogenizing in ice-cold RIPA buffer (Thermofisher, 89900). Homogenates were centrifuged at 15,000 g for 15 min at 4°C. The resulting supernatants were collected, and protein content determined by Quick Start Bradford kit assay (Bio-Rad, 500-0203). 20-50 μg of total protein was electrophoresed on 10% SDS– polyacrylamide gel, transferred to a nitrocellulose blotting membrane (Amersham Protran 0.45 μm, GE Healthcare Life Sciences, 10600002) and blocked in TBS-T (150 mM NaCl, 20 mM Tris–HCl, pH 7.5, 0.1% Tween 20) supplemented with 5% non-fat dry milk. Membranes were incubated overnight at 4°C with the OCT4 primary antibody (Oct-3-4, C-10, Santacruz, sc-5279) in TBS-T supplemented with 5% non-fat dry milk, washed with TBS-T and next incubated with secondary HRP-conjugated anti-mouse IgG (1:2,000, DAKO, P0447) for 1 hour at room temperature and developed using the ECL detection kit (Perkin Elmer, NEL105001EA).

### Data analysis

Statistical analysis was performed using SPSS 21.0 (SPSS® Statistic IBM®). The normality of the data was studied by Shapiro-Wilk test and homogeneity of variance by Levene test. For comparison of two independent groups, two-tail unpaired t-Student’s test (data with normal distribution), Mann-Whitney-Wilcoxon or Kolmogorov-Smirnov tests (with non-normal distribution) was executed. For multiple comparisons, data with a normal distribution were analyzed by one way-ANOVA test followed by a Tukey’s (equal variances) or a Games-Howell’s (not assumed equal variances) post-hoc tests. Statistical significance of non-parametric data for multiple comparisons was determined by Kruskal-Wallis one-way ANOVA test.

## SUPPLEMENTAL FIGURE LEGENDS

**Figure S1:**
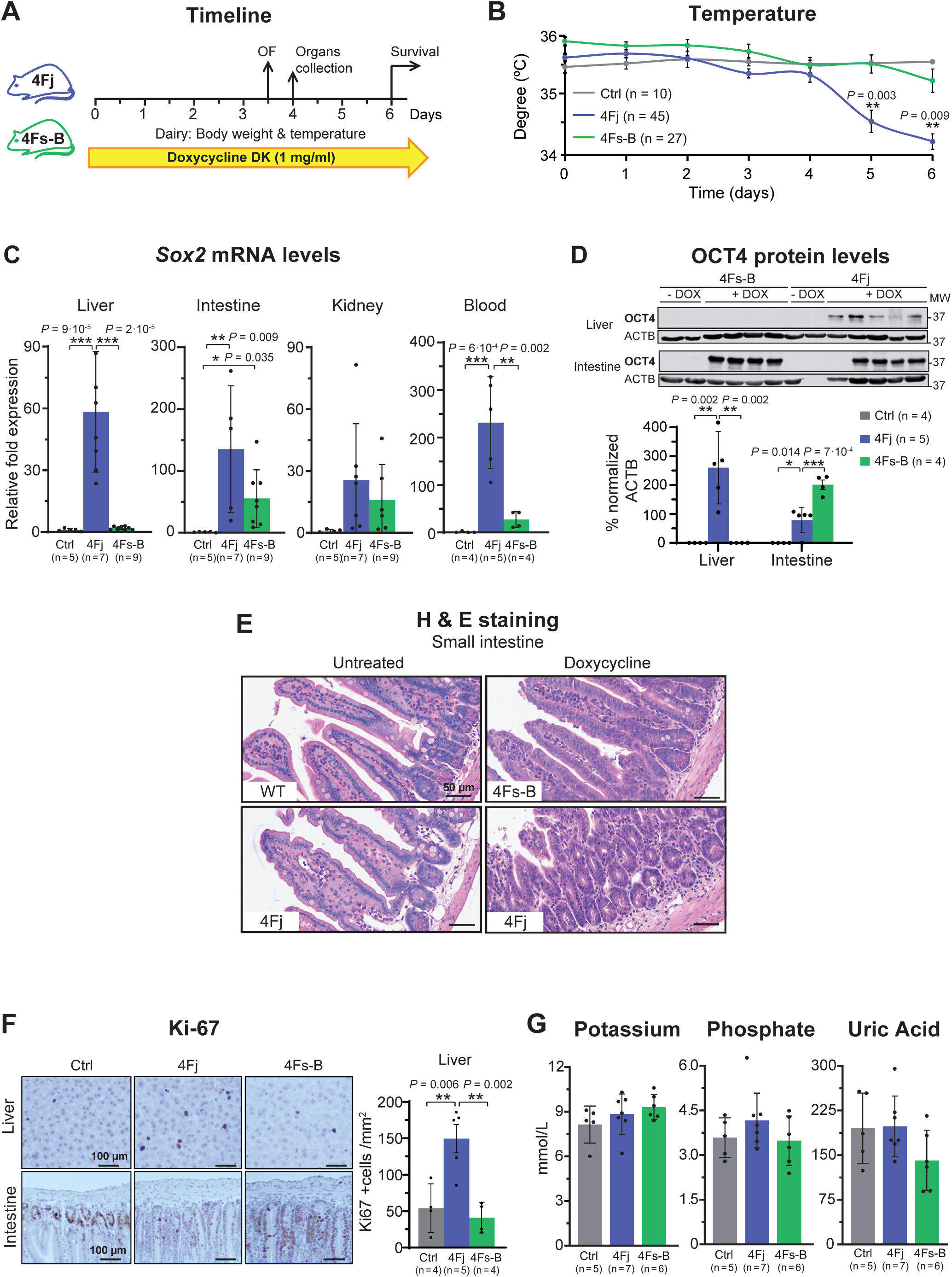
In vivo reprogramming induces liver and intestinal damage. (A) Experimental design indicating doxycycline administration protocol, activity measure by open field (OF) test, and organs and plasma collection. (B) Superficial temperature of 4Fj and 4Fs-B mice upon continuous administration of doxycycline, as wells as untreated 4F control mice (Ctrl). (C) *Sox2* mRNA and (D) OCT4 protein levels in the indicated organs of untreated 4F controls, and 4Fj and 4Fs-B mice after induction of 4 days. (E) Small intestine hematoxylin and eosin staining for an untreated wildtype and 4Fj mice and treated 4Fj and 4Fs-B mice during 4 days. (F) Immunostaining and quantification of Ki67 positive cells in the liver and small intestine (n = 3 regions from four untreated controls, five 4Fj, and four 4Fs-B mice). Scale bars, 100 μm. (G) Potassium, phosphate, and uric acid levels in plasma of untreated control mice (Ctrl), and 4Fj and 4Fs-B mice after 4 days of induction of in vivo reprogramming. Data are mean ± SEM. Statistical significance was assessed by (B-D, F-G)) one-way ANOVA followed by Tukey’s or Games-Howells post hoc tests.

**Figure S2:**
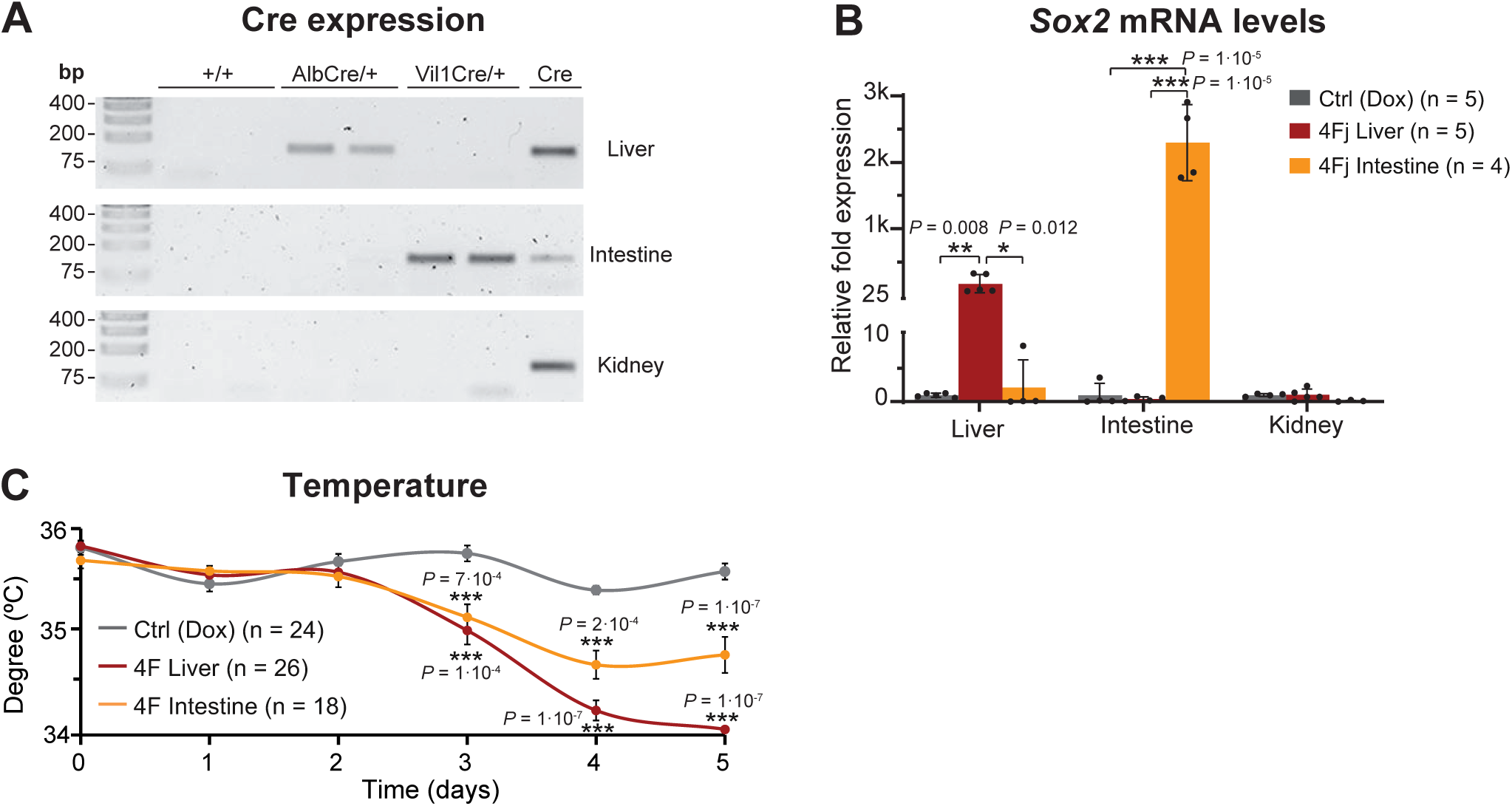
Liver and intestine-specific reprogramming lead to decrease body weight and early mortality. (A) Expression of Cre recombinase in the liver, small intestine, and kidney of control (4Fj LSL-rtTA), 4F-Liver (4Fj LSLrtTA AlbCre), and 4F-Intestine (4Fj LSLrtTA Vil1Cre) mice. (B) *Sox2* mRNA levels in control, 4F-Liver, and 4F-Intestine mice after induction of reprogramming for 4 days. (C) Superficial body temperature upon continuous administration of doxycycline in controls and 4F tissue-specific reprogramming mice. Data are mean ± SEM. Statistical significance was assessed by (B-C) one-way ANOVA followed by Tukey’s post hoc test.

**Figure S3:**
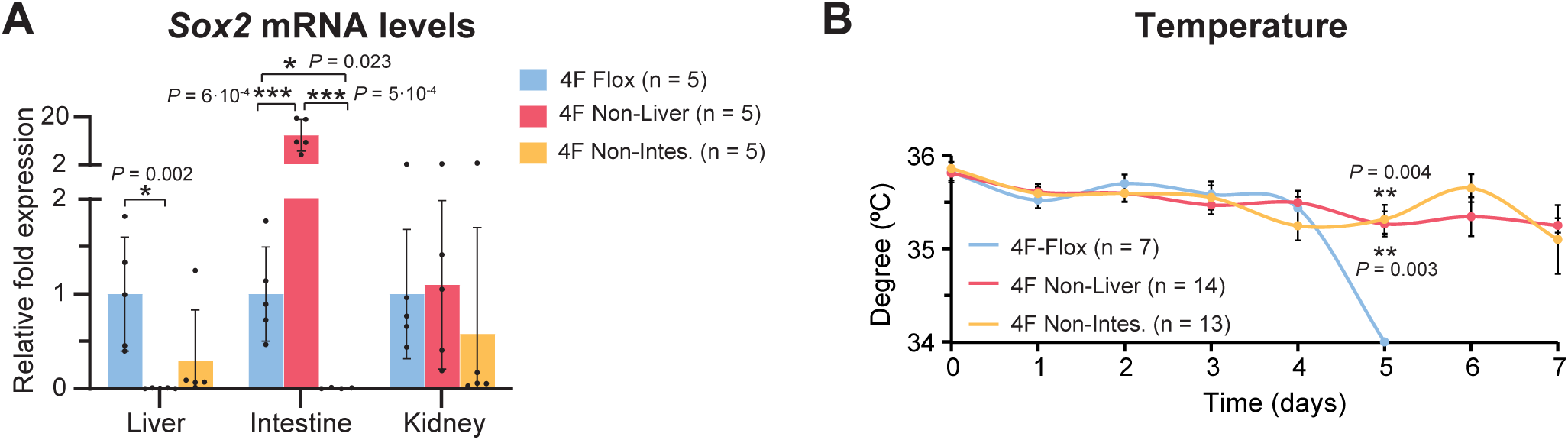
Avoiding the expression of OSKM in liver and intestine diminishes adverse phenotypes and premature death associated with in vivo reprogramming. (A) *Sox2* mRNA levels in in the liver, small intestine, and kidney of 4F-Flox (OSKM whole body), 4F Non-Liver, and 4F Non-Intestine mice, after 4 days of induction of in vivo reprogramming. (B) Superficial body temperature upon continuous administration of doxycycline in whole body (4F-Flox), 4F Non-liver, and 4F Non-intestine mice. Data are mean ± SEM. Statistical significance was assessed by (A-B) one-way ANOVA followed by Tukey’s post hoc test.

**Figure S4:**
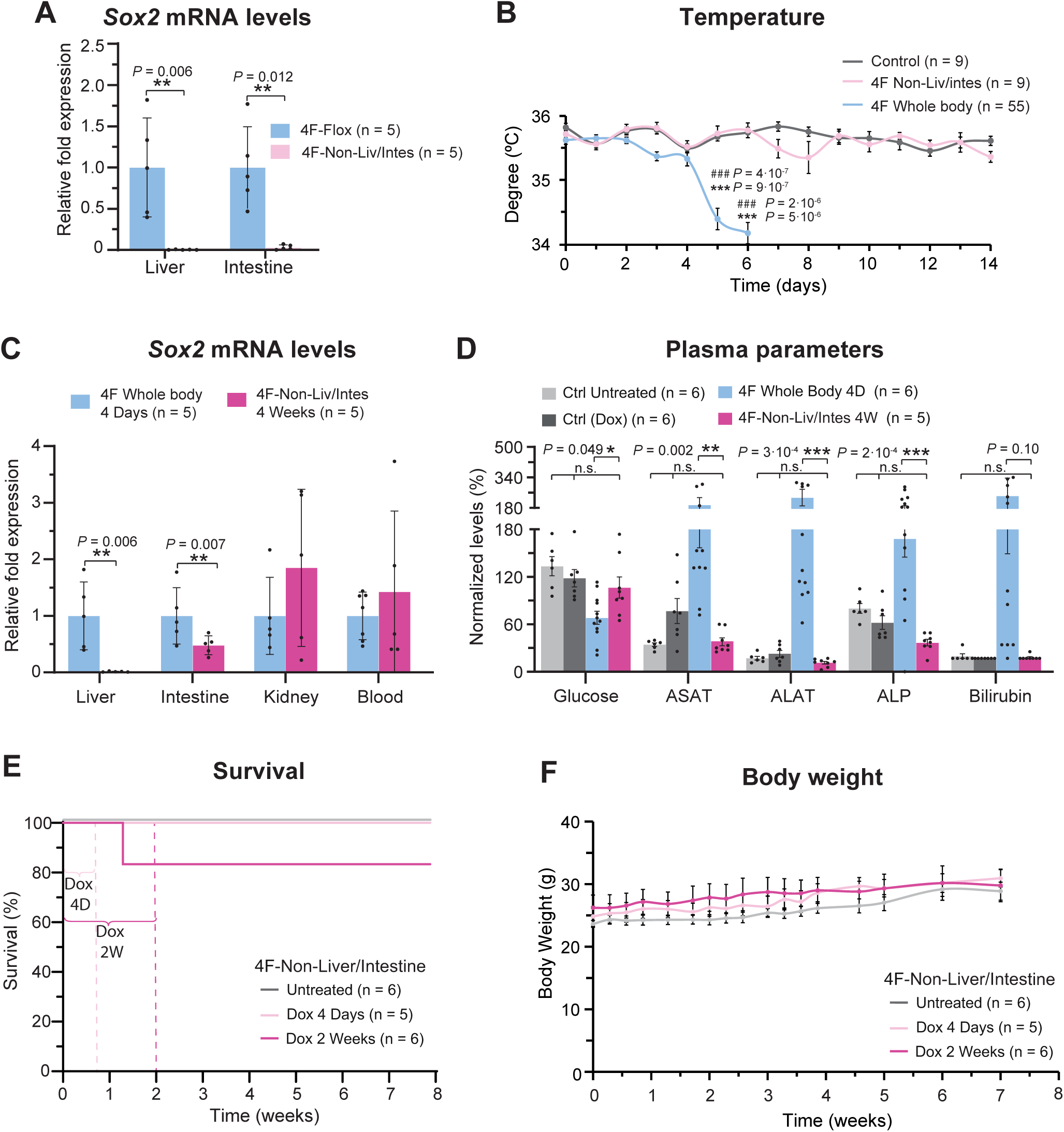
Safe and long-term continuous induction of in vivo reprogramming in a novel reprogrammable strain. (A) *Sox2* mRNA levels in control and 4F non-Liver/Intestine mice after induction of in vivo reprogramming for 4 days. (B) Body temperature of control and 4F Non-Liver/Intestine mice upon continuous administration of doxycycline. (C) mRNA levels of *Sox2* in multiple organs of 4F whole body reprogrammable and 4F Non-Liver/Intestine mice after 4 days and 4 weeks of continuous doxycycline treatment. (D) Glucose, liver enzymes (ASAT, ALAT, ALP) and bilirubin levels in plasma of untreated 4F controls, treated no reprogrammable mice, 4F whole body after 4 days of treatment, and 4F Non-Liver/Intestine mice after 4 weeks of induction of in vivo reprogramming. (E) Survival and (F) body weight of 4F Non-Liver/Intestine mice non-treated or doxycycline-treated for 4 days or 2 weeks and monitored after doxycycline withdrawal. Data are mean ± SEM. (A, C) Two-sided unpaired t-test and Mann-Whitney-Wilcoxon test, (B, D, F) one-way ANOVA followed by Tukey’s post hoc test and (E) log-rank (Mantel-Cox) test.

## AUTHOR CONTRIBUTIONS

A.P. was involved in the design of the study, in all experiments, data collection, and statistical analysis. A.V. performed mouse doxycycline treatment, immunostainings, and qRT-PCR experiments. G.D. and C. M. contributed to RNA and protein extraction, qRT– PCR, and western blot analysis. K.P. made intellectual contributions. C.R. and F.B. was implicated in RNA extraction and qRT-PCR experiments. C.B. was involved in mouse genotyping and sample collection. A.O. directed and supervised the study and designed the experiments. A.P. and A.O wrote the manuscript with input from all authors.

## ACKNOWLEDGMENTS

We would like to thank all members of the Ocampo laboratory for helpful discussions. We would like to thank the teams of mouse facilities at the University of Lausanne including Francis Derouet (head of the animal facility at Epalinges), Lisa Arlandi, Isabelle Grandjean (head of the animal facility of Agora) and Laurent Lecomte (head of the animal facility of the Department of Biomedical Sciences). We thank Andrea Ablasser for the kind donation of the 4Fs-B rtTA mice. We thank the Service de Chimie Clinique du CHUV for mouse plasma analysis and the Mouse Pathology Facility of the University of Lausanne for tissue processing and immunostainings.

